# Dynamics of the MRSA Population in A Chilean Hospital: A Phylogenomic Analysis (2000-2016)

**DOI:** 10.1101/2023.02.06.526811

**Authors:** José RW Martínez, Paul J. Planet, Spencer-Sandino Maria, Rivas Lina, Díaz Lorena, Quesille-Villalobos Ana, Riquelme-Neira Roberto, Alcalde-Rico Manuel, Hanson Blake, Lina P Carvajal, Rincón Sandra, Reyes Jinnethe, Lam Marusella, Araos Rafael, García Patricia, César A. Arias, José M. Munita

**Affiliations:** Genomics & Resistant Microbes (GeRM), ICIM, Facultad de Medicina Clínica Alemana, Universidad del Desarrollo, Chile; Multidisciplinary Initiative for Collaborative Research On Bacterial Resistance (MICROB-R); Division of Pediatric Infectious Diseases, Children’s Hospital of Philadelphia, Philadelphia, USA; Department of Pediatrics, Univ. of Pennsylvania, Philadelphia, USA; American Museum of Natural History, New York, USA; Molecular Genetics and Antimicrobial Resistance Unit, Universidad El Bosque, Colombia; Núcleo de Investigaciones Aplicadas en Ciencias Veterinarias y Agronómicas, Facultad de Medicina Veterinaria y Agronomía, Universidad de Las Américas, Chile; Grupo de Resistencia a los Antibióticos en Bacterias Patógenas y Ambientales (GRABPA), Pontificia Univ. Católica de Valparaíso; Center for Antimicrobial Resistance and Microbial Genomics, Univ of Texas Health Science Center, McGovern Medical School, Houston, Texas, USA; Departamento de Laboratorios Clínicos, Escuela de Medicina, Pontificia Universidad Católica de Chile; Division of Infectious Diseases, Houston Methodist Hospital, Houston, USA; Center for Infectious Diseases, Houston Methodist Research Institution, Houston, TX, USA; Department of Medicine, Weill Cornell Medical College, New York, USA; Hospital Padre Hurtado, Chile

## Abstract

The global dissemination of methicillin-resistant *Staphylococcus aureus* (MRSA) is associated with the emergence and establishment of clones in specific geographic areas. The Chilean-Cordobes clone (ChC) (ST5-SCC*mec*I) has been the predominant MRSA clone in Chile since its first description in 1998, despite the report of other emerging MRSA clones in the last years. Here, we characterize the evolutionary history of MRSA from 2000 to 2016 in a Chilean tertiary healthcare center using phylogenomic analyses. We sequenced 469 MRSA isolates collected between 2000-2016 in a tertiary healthcare center in Chile. We evaluated the temporal trends of the circulating clones and performed a phylogenomic reconstruction to characterize the clonal dynamics. We found a significant increase in the diversity and richness of sequence types (STs; Spearman r=0.8748, p<0.0001) with a Shannon diversity index increasing from 0.221 in the year 2000 to 1.33 in 2016. The temporal trend analysis revealed that in the period 2000-2003 most of the isolates (94.2%; n=98) belonged to the ChC clone. However, since then, the frequency of the ChC clone has decreased over time, accounting for 52% of the collection in the 2013-2016 period. This decline was accompanied by the rise of two emerging MRSA lineages, ST105-SCC*mec*II and ST72-SCC*mec*VI. In conclusion, the ChC clone remains the most frequent MRSA lineage in Chile. However, this lineage is gradually being replaced by several emerging clones, the most important of which is clone ST105-SCC*mec*II. To the best of our knowledge, this is the largest study of MRSA clonal dynamics performed in South America.

**Importance:** Methicillin-resistant *Staphylococcus aureus* (MRSA) is a major public health pathogen that disseminates through the emergence of successful dominant clones in specific geographic regions. Knowledge of the dissemination and molecular epidemiology of MRSA in Latin America is scarce and is largely based on small studies or classical typing techniques with several limitations to depict an accurate description of their genomic landscape. We used whole-genome sequencing to study 469 MRSA isolates collected between 2000-2016 in Chile to provide the largest and most detailed study of clonal dynamics of MRSA carried out in South America to date. We found a significant increase in the diversity of MRSA clones circulating over the 17-year study period. Additionally, we describe the emergence of two novel clones (ST105-SCCmecII and ST72-SCCmecVI), which have been gradually increasing their frequency over time. Our results drastically improve our understanding of the dissemination and update our knowledge about MRSA in Latin America.

## Introduction

Infections caused by methicillin-resistant *Staphylococcus aureus* (MRSA) are a major public health problem. The World Health Organization (WHO) has listed MRSA as one of the high-priority pathogens to target research and development of new antimicrobials (1). MRSA spread occurs largely through the successful dissemination of specific genomic lineages in certain geographic regions (2). Clonal replacement waves characterized by the emergence of new MRSA clones with greater adaptive capacity have been well documented (2,3). However, knowledge about the factors that drive the clonal replacement phenomenon in a specific geographical region is scarce. Improving our understanding of this phenomenon is crucial to design strategies to contain MRSA spread.

A key aspect in the surveillance of MRSA is the molecular characterization of circulating lineages to identify outbreaks and sources of colonization, distinguish between community and hospital strains, and inform infection control strategies (4–6). MRSA lineages have been classically described using molecular typing methods such as pulsed-field gel electrophoresis (PFGE), multi-locus sequence typing (MLST), and characterization of the type of *SCCmec* cassette or *spa* gene (7–10). However, current whole-genome sequencing (WGS)-based analyzes provide greater granularity in the description of the evolution of bacterial organisms and higher resolution to differentiate specific genomic lineages within strains classified as part of the same clonal group by classic typing methods (11–14). Therefore, the use of WGS provides the means for a better description and understanding of the evolutionary history of human pathogens such as MRSA.

Detailed characterizations of endemic MRSA clones in low- and middle-income countries are scarce, with most of the data coming from developed regions (15–17). A prospective study including MRSA isolates recovered from the bloodstream of six Latin American countries between 2011-2013 reported that the major MRSA clones circulating in the region belonged to the clonal complex (CC) 5, CC8, and CC30, with major differences among countries (18). The same study found >90% of MRSA circulating in Chile and Peru belonged to the Chilean-Cordobes (ChC) clone, an ST5-SCC*mec*I lineage first described in Chile and Argentina in 1998 (18,19). This ChC clone rapidly spread throughout Latin America in the early 2000s, displacing other dominant clones (20–22). However, it was later almost entirely replaced in Ecuador, Colombia, and Venezuela by a community-associated ST8-SCC*mec*IV lineage designated USA300 Latin American Variant (USA300-LV) (20,23–25). Interestingly, the few available data, all based on classical typing techniques, suggest this clonal replacement was not observed in Chile and Perú, where the ChC clone remains predominant, despite the well-reported circulation of successful pandemic lineages such as USA300 and USA300-LV (6,18). The factors underlying this differential behavior of clonal MRSA spread within countries of the same geographical region remain largely unknown.

In this study, we aimed to provide a detailed picture of the population dynamics of MRSA clones in Chile using a collection of 469 MRSA isolates sequentially obtained between 2000-2016 in a tertiary healthcare center located in Santiago, Chile. We report for the first time an ongoing gradual clonal replacement of the ST5-SCC*mec*I ChC clone by emerging MRSA lineages that include ST1*05-SCCmec*II and ST72-SCC*mec*IV-VI. Our data constitute the largest and most detailed study of clonal dynamics of MRSA carried out in South America to date.

## Materials and methods

### Collection of MRSA strains

We included a total of 469 MRSA isolates collected between 2000-2016 in the Hospital Clinico UC-Christus in Santiago, Chile. All isolates were identified as *S. aureus* using MALDI-TOF (Bruker). Methicillin resistance was confirmed with a cefoxitin disk diffusion assay, following the recommendations of the Clinical and Laboratory Standards Institute (CLSI) (26). The *mecA* gene was detected by PCR assays as previously described (27).

### Whole-genome sequencing and genomic characterization

Genomic DNA of all 469 isolates was purified with the DNeasy^®^ Blood & Tissue kit (Qiagen) from fresh cultures treated with lysostaphin for 30 minutes at 37°C. DNA concentration was determined by Qubit™ dsDNA HS assay on a Qubit 2.0 fluorometer (Thermo Fisher Scientific). Genomic libraries were prepared with the NexteraXT DNA Sample Preparation Kit (Illumina) and WGS was performed using Illumina MiSeq and HiSeq instruments, generating paired reads of 150 or 300 nucleotides. The quality of the raw reads was determined by FASTQC v0.11.9 and MultiQC v1.10.1 (https://www.bioinformatics.babraham.ac.uk/projects/fastqc/) (28). Raw reads were trimmed with Trimmomatic v0.39; those reads exhibiting a Phred quality score < 30 were excluded (29). Finally, the genomes were *de novo* assembled with SPAdes v3.13.0 and the quality of the assemblies was evaluated with QUAST v5.0.2 (30,31). *In silico* characterization of the ST and *SCCmec* types was performed using MLST v2.19.0 (https://github.com/tseemann/mlst) and staphopia-sccmec, respectively (32,33). In addition, the presence of antimicrobial resistance genes, virulence factors, and genetic traits associated with resistance to heavy metals was assessed with ABRicate v1.0.1 (https://github.com/tseemann/abricate).

### Phylogenomic analyzes

Genomes were annotated with Prokka v1.14.5 and a multiple alignment with a core-genome definition of 99% was performed with Roary v3.13.0 (34,35). RAxML 8.2.12 was used to construct a maximum likelihood (ML) phylogenomic tree using a GTR Substitution Model with a bootstrap of 100 to then generate a recombination-free ML tree with ClonalFrameML v1.12 (36,37). The phylogenomic trees were visualized with the Tree Of Life (iTOL) v6 interactive tool (42).

### Statistical analysis

The normal distribution of the data was evaluated with the D’Agostino & Pearson normality test. Diversity parameters were determined with the Shannon diversity index using the Omni calculator tool (https://www.omnicalculator.com/). The correlation between ST diversity and isolation year was determined by the Spearman ranked test. For the analysis of temporal dynamics, genomes were grouped into 4 time periods according to their year of isolation to obtain a similar number of genomes in each period: 2000-2003, 2004-2008, 2009-2012, and 2013-2016.

## Results

### The diversity of circulating MRSA lineages significantly increased over time

Of a total of 469 MRSA isolates, 192 (41%) were recovered from the respiratory tract, 173 (36.9%) from the bloodstream or other sterile sites, 94 (20%) from the skin, and 10 (0.2%) from urine samples (Table 1). Our genomic analyzes revealed a total of 15 different sequence types (ST) among the 469 MRSA genomes, with 91% of the isolates belonging to only three STs; ST5 (n=329, 70.1%), ST105 (n=70, 14.9%), and ST72 (n=28, 6%) (Table 1). To evaluate the changes of STs over time, we performed a temporal analysis grouping the genomes in 4 time periods: 2000-2003, 2004-2008, 2009-2012, and 2013-2016. This analysis demonstrated a significant increase in the diversity of STs over the years (Spearman r = 0.8748, p<0.0001). Indeed, the Shannon diversity index for STs increased from 0.2 in the 2000-2003 period, to 1.4 in the 2013-2016 period (Figure 1). Furthermore, while MRSA isolates collected during 2000-2003 belonged to only two different STs (ST5 and ST239), the genomes from the 2013-2016 period encompassed 13 different STs (Figure 2 and S1). Interestingly, as shown in Figure S1, the increase in ST diversity was observed in all anatomical sites, except for the bloodstream, where we only found four STs (ST5, ST105, ST72, and ST239) throughout the 17-year study period (Figure S1).

**Table 1.**
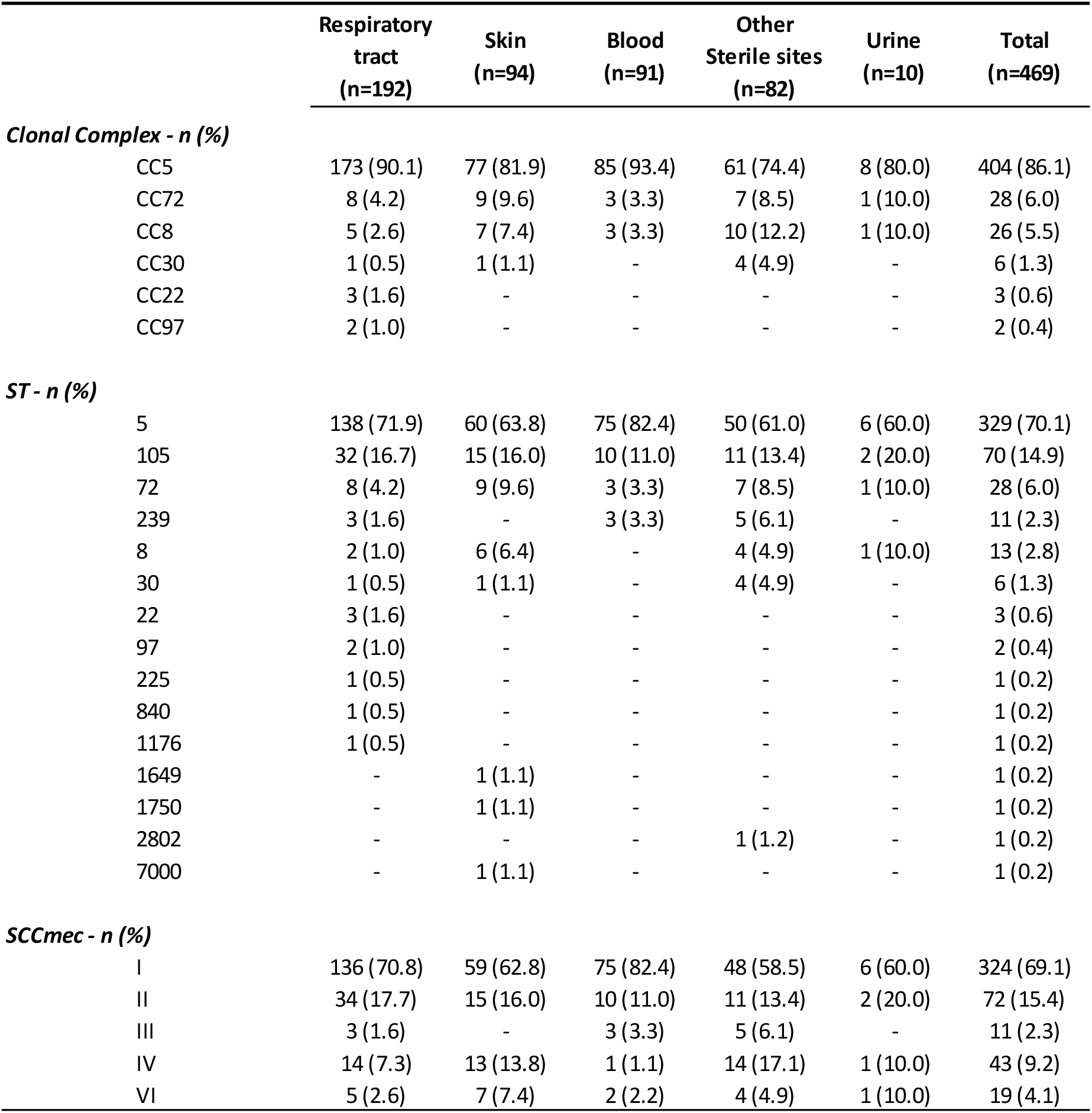
Distribution of Clonal complex, ST and SCC*mec* by anatomical origin.

**Figure 1.**
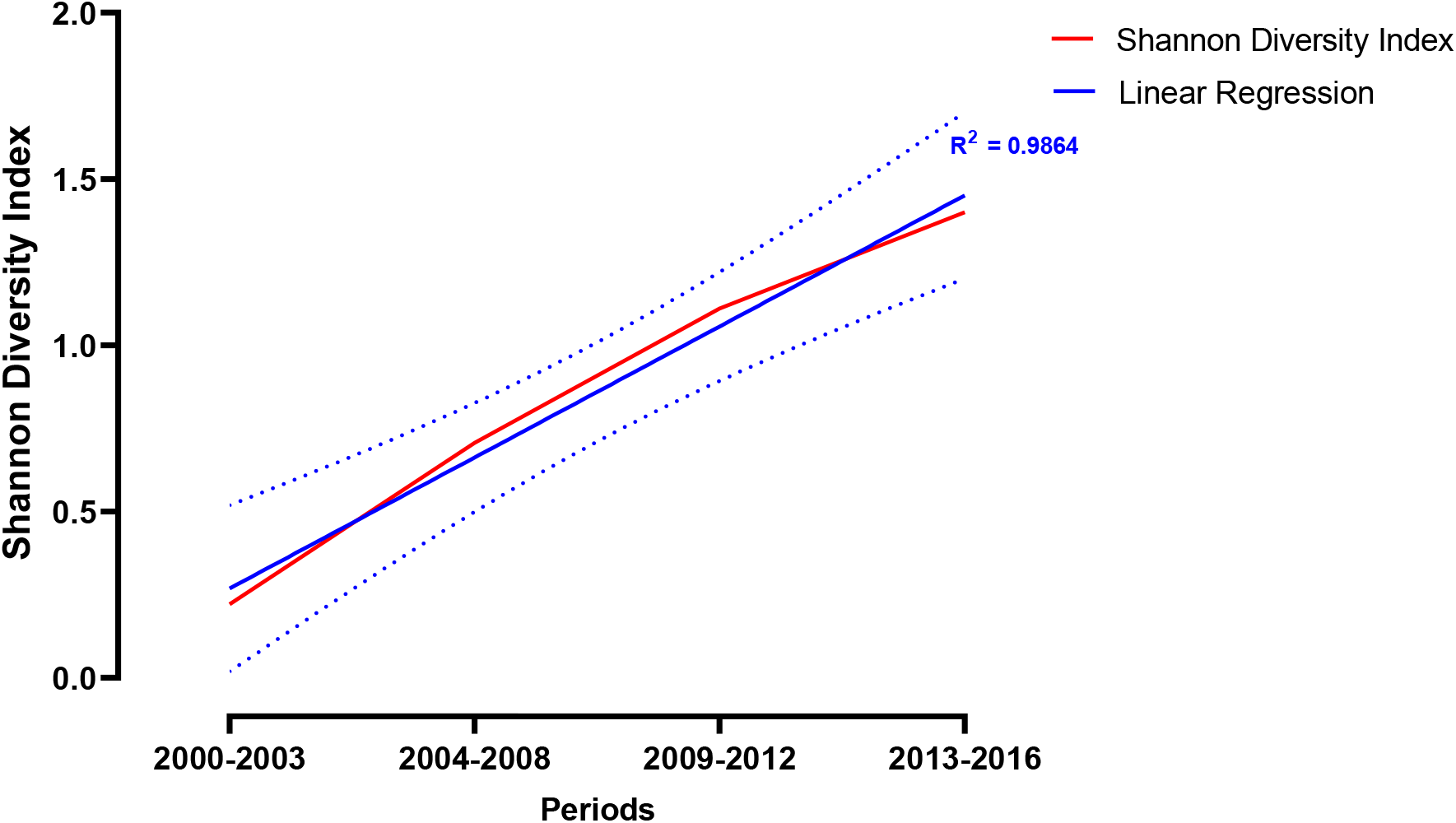
Diversity of STs over time. The axes indicate the Shannon diversity index (Y-axis) and the year of isolation (X-axis). The blue line represents the Shannon diversity index for each year. The continuum blue line shows the linear regression and the dotted blue line the 95% confidence interval.

**Figure 2.**
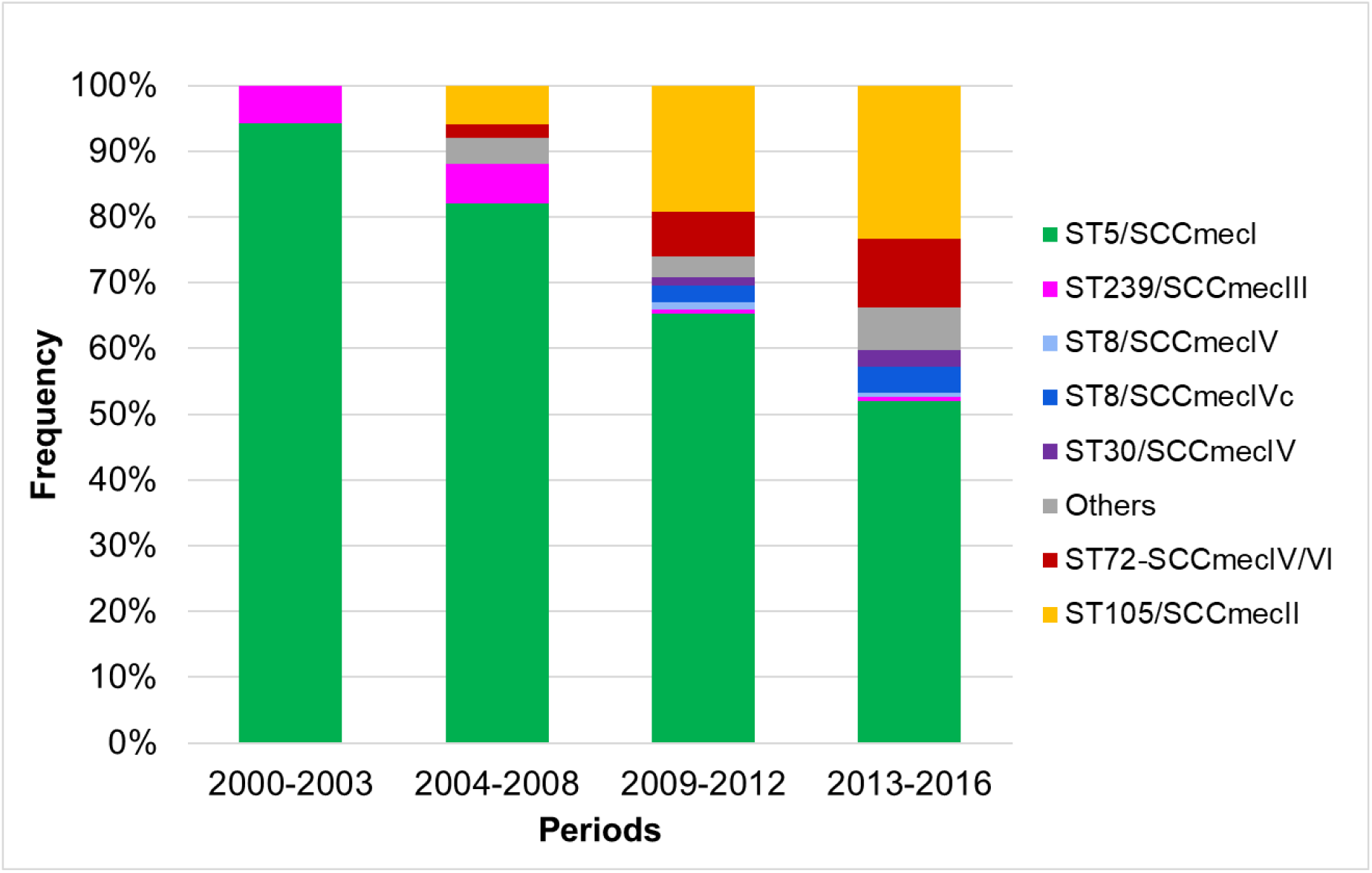
Temporal trends of MRSA clones. Stacked bar plot showing the frequency (% of isolates) of the different clones identified in each of the periods.

### Gradual replacement of the ChC ST5 MRSA clone and emergence of ST105 and ST72 lineages over the years

To further characterize the clonal dynamics of MRSA, we used the combination of ST and SCC*mec* types and evaluated their frequency over the four aforementioned time periods. Our temporal trend analysis revealed most isolates obtained during the first period (2000-2003) belonged to the ChC ST5-SCC*mec*I lineage (n=98, 94.2%), and the few remaining corresponded to ST239-SCC*mec*III (n=6, 5.8%) (Figure 2). Interestingly, a gradual reduction in the frequency of the ChC clone over time was observed, dropping from the initial 94.2% to only 52% in the 2013-2016 period (Figure 2). In parallel, an increase of two emerging MRSA lineages was observed: ST105 and ST72, neither of which were circulating in the first period of observation (Figure 2 and S1). All ST105 genomes carried a SCC*mec*II type (ST105-SCC*mec*II). The frequency of ST105-SCC*mec*II gradually increased over time from 0% in 2000-2003 to 6% in 2004-2008, 19.3% in 2009-2012, and 23.4% in the 2013-2016 period (Figure 2 and S1). In the case of ST72, two types of *SCCmec* were observed (ST72-SCC*mec*IV and ST72-SCC*mec*VI). Similar to ST105, a gradual increase was observed in ST72 frequency, from not detected in 2000-2003 to 2.0%, 6.8%, and 10.4% in the 2004-2008, 2009-2012, and 2013-2016 periods, respectively (Figure 2 and S1).

We then evaluated the temporal trends of clonal frequency by anatomical site. While the frequency of the ChC ST5-SCC*mec*I clone decreased in all sites, this reduction was particularly marked in samples collected from the skin and bloodstream, with frequencies decreasing from 100% (n=19) and 93.5% (n=43) in the 2000-2003 period to 39.4% (n=13) in the skin and 37.5% (n=3) in the bloodstream during the 2013-2016 period (Figure S1). On the other hand, while the frequency of ST105-SCC*mec*II clone increased proportionally across all anatomical sites, the ST72SCC*mec*IV/VI lineages presented a higher frequency in isolates from the skin (21.2%) and bloodstream (25%), and lower frequency in respiratory samples (4.3%) during the 2013-2016 period (Figure S1).

### Genomic characterization of the emerging ST105 and ST72 MRSA clones

We performed a core-genome-based phylogenomic reconstruction to understand the genomic relationships among MRSA lineages. Additionally, to further analyze relevant characteristics within MRSA lineages circulating over this 16-year period, we also searched for antimicrobial resistance genes, virulence factors, and heavy metal resistance genes. Our phylogenomic reconstruction revealed six well-defined clades (Figure 3). Clade-1 was only composed of six ST30-SCC*mec*IVc genomes (CC30). Clade-2 consisted of three ST22-SCC*mec*IVc genomes and clade-3 included only two ST97-SCC*mec*IVa genomes. Clade-4 grouped 26 CC8 genomes: 11 ST239-SCC*mec*III; 11 ST8-SCC*mec*IVc; two ST8-SCC*mec*IVa; one ST2802-SCC*mec*IVc and one ST1750-SCC*mec*IVa. All 28 genomes from clade-5 belonged to the ST72 lineage, of which 19 carried SCC*mec*VI and 9 SCC*mec*IVc. Finally, clade-6 grouped most of the isolates (n=403), all of which belonged to CC5. Of them, 325 were ST5-SCC*mec*I (ChC clone); 70 were ST105-SCC*mec*II; two ST5-SCC*mec*IVc and the remaining 6 corresponded to one of each of the following: ST5-SCC*mec*IVa, ST5-SCC*mec*II, ST225-SCC*mec*II, ST840-SCC*mec*IVc, ST1176-SCC*mec*IVa, and ST7000-SCC*mec*IVc.

**Figure 3.**
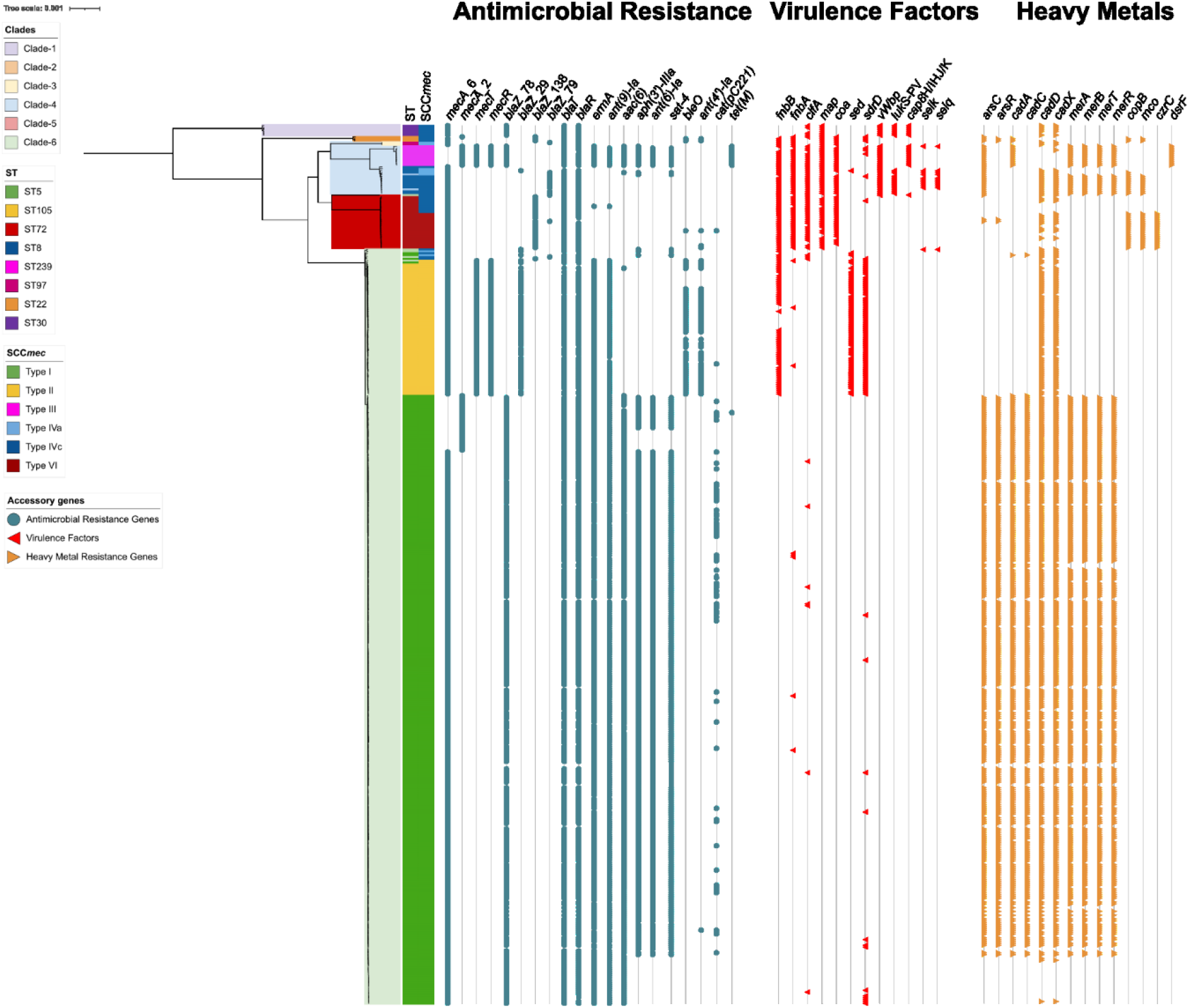
Phylogenomic reconstruction of the 469 isolates of MRSA. Phylogenomic reconstruction rooted to the midpoint of genomic distances. The most important clades are represented by colors within the reconstruction. The colored bands indicate the genetic lineage (ST and SCC*mec*). Dots show the presence of antimicrobial resistance genes. Red triangles indicate the presence of virulence factors. Yellow triangles indicate the presence of heavy metal resistance genes.

All genomes harbored *mecA* and >99% carried the antimicrobial resistance genes, virulence factors, and heavy metal resistance genes detailed in the supplementary Table 1. Interestingly, despite belonging to the same clonal complex (CC5), the emerging ST105-SCC*mec*II lineage exhibited important differences with the ST5-SCC*mec*I ChC clone (Figure 3). The genomes of ST105-SCC*mec*II lacked the aminoglycoside resistance genes *AAC(6’)-Ie-APH(2”)-Ia*, and *ant(6)-Ia*, and the erythromycin and quinolone resistance gene *sat-4*, all of which were frequently harbored by isolates of the ChC clone (97.8%; 86.5%; and 87.1%, respectively) (Figure 3). In contrast, the ST105-SCC*mec*II lineage frequently carried the aminoglycoside resistance gene *ant(4)-Ia* (75.7%), the bleomycin resistance gene *bleO* (75.7%); and the virulence factors *sdrD* (97.1%), related to immune evasion, the enterotoxin *sed* (90%), and *fnbB* (80%), which encodes a fibrinogen binding protein associated with biofilm formation. Notably, all of these genetic traits were absent in the ST5-SCC*mec*I ChC clone (Figure 3). In addition, while a large proportion of the ChC clone presented the heavy metal resistance clusters *arsBCR* (81.8%), *cadAC* (81.5%), and *merABTR* (79.4%), these genetic traits were absent in genomes of the ST105-SCC*mec*II lineage. Both ST5-SCC*mec*I and ST105-SCC*mec*II frequently carried the *cadDX* cadmium resistance genes (91.4% and 81.5%, respectively) (Figure 3).

Similarly, the emerging ST72 lineage lacked several of the antimicrobial resistance genes present in the ChC clone but displayed a high frequency of virulence factors such as *fnbA* (93%), *fnbB* (100%), *clfA* (89%), *coa* (89%) and *map* (82%) (Figure 3). Interestingly, the main difference between the ST72-SCC*mec*IV and ST72-SCC*mec*VI was the presence of the heavy metal resistance genes *copB, mco*, and *czrC* in all the ST72-SCC*mec*VI isolates, none of which were observed in ST72-SCC*mec*IV (Figure 3).

## Discussion

In this study, we performed WGS on a collection of 469 invasive MRSA isolates obtained in a tertiary healthcare center from Santiago, Chile, over a 17-year period (2000-2016) with the aim of providing a detailed picture of the clonal dynamics of MRSA. Our data suggest that there was a significant increase in the diversity of circulating MRSA clones over the years. In addition, we report an ongoing gradual clonal replacement of the ST5-SCC*mec*I ChC clone by two emerging MRSA lineages that include ST105-SCC*mec*II and ST72-SCC*mec*IV-VI.

Our first relevant observation was an important increase in the diversity of STs of MRSA circulating during the study period. Indeed, while only two STs were found during 2000-2003, this number increased to 13 during the subsequent periods of observation. Our findings differ from the classical description of clonal MRSA replacement events, where the number of circulating clones is limited, and dominant lineages are replaced by new, better-adapted clones (2). However, a recent multicenter study performed in China over a 6-year period (2014-2020) reported an increase in the diversity of circulating MRSA lineages over time (43). A key common aspect between this study and our current report is the use of WGS-based analyses, which likely results in better resolution to differentiate closely related isolates that could otherwise be classified as the same clone by classical typing techniques.

Our data confirm the ST5-SCC*mec*I ChC clone continues to be the most predominant MRSA lineage in the country. However, our findings also suggest the ChC clone is gradually being replaced by two emerging MRSA lineages, ST105 and ST72. The ST105-SCC*mec*II lineage is closely related to the ChC clone as they both belong to the same clonal complex (CC5). Interestingly, a recent study described the emergence of an ST105-SCC*mec*II lineage (designated as RdJ clone) between 2014-2017 in Rio de Janeiro, Brazil, showing a strong association with bloodstream infections and with a higher rate of immune evasion (44). In line with these findings, our analysis stratifying by the source of isolation demonstrated that the ST105-SCC*mec*II lineage first emerged in bloodstream samples (Figure S1). Also, our data suggest the magnitude of the clonal replacement of the ChC clone differs among anatomical sites, with higher prevalence in respiratory samples as opposed to blood or skin (Figure S1). The factors underlying these differences are not fully understood and require further studies. It is also worth mentioning that Arias *et al*. also described the ST105-SCC*mec*II lineage in Chile and Brazil, in 2013 (18). However, genomic analyses revealed important differences between isolates belonging to this lineage recovered in these two countries. In particular, ST105-SCC*mec*II isolates from Brazil harbored the aminoglycoside resistance gene *APH(3’)-III*, and the macrolide resistance genes *msrA* and *mphC*, none of which were found in Chilean isolates (18). Similarly, genomes of ST105-SCC*mec*II from our collection lacked *APH(3’)-III, msrA, mphC*, and *chp*,suggesting that the emerging clone observed herein belongs to the same lineage previously described by Arias and colleagues (18). Altogether, our observations suggest that despite being part of the same lineage, ST105-SCC*mec*II isolates emerging in Chile and Brazil exhibit potentially important genomic differences. The study of the implications of these disparities is part of our ongoing research efforts.

The other emerging lineages observed in our results belong to ST72 and include the ST72-SCC*mec*IV and ST72-SCC*mec*VI. The former is a genetic lineage previously described in community-associated infections in South Korea (45–47). In contrast, to the best of our knowledge, this is the first description of ST72-SCC*mec*VI. Our phylogenomic analysis suggested these two clones are closely related, as observed in Figure 3. Interestingly, previous studies have included ST72-SCC*mec*IV as part of the CC8 lineage (48–50). However, our phylogenomic analysis revealed that ST72 forms a strongly distinct clade from other CC8 genomes. Indeed, the branch length distances between ST72 (SCC*mec*IV and SCC*mec*VI) and the CC8 genomes were higher than between the genomes of the CC96 and the CC8, suggesting that the ST72 clone should be included in a new and independent clonal complex. More studies are needed to perform a deeper characterization of the new emerging clone ST72-SCC*mec*VI described herein.

Our study has limitations worth mentioning. First, our isolate collection was limited to one healthcare center, therefore preventing us from extrapolating our results about the dynamics of MRSA clonal replacement to a country-wide or regional level. However, we expect our data to shed light on the molecular epidemiology of MRSA in Latin America and help future studies to compare the situation in other countries in the region. Second, we did not obtain access to the clinical metadata, hampering our ability to explore possible associations between specific MRSA lineages with clinical presentations and patient outcomes. Finally, although all MRSA isolates were obtained from hospitalized patients, our data did not allow us to accurately discriminate between hospital and community-acquired infections, which would have allowed us to better understand the role different MRSA lineages are playing in these clinical presentations. A more complete study including isolates from different geographic areas in a broad spanning time with their associated clinical metadata will be helpful to obtain a better picture of the genomic landscape of the MRSA clonal dynamics in Latin America and its clinical implications.

In summary, we have characterized an atypical clonal replacement phenomenon, demonstrating the usefulness of WGS in the study of pathogens of a major public health problem such as MRSA. Although this study includes isolates only from a single healthcare center, it is the most robust and detailed study of the genomic epidemiology of MRSA conducted in the country to date. Additionally, this study highlights a cryptic clonal replacement phenomenon where the ChC clone is gradually being replaced by several genetic lineages, a phenomenon that differs from that classically described for MRSA. The results obtained here will allow a better understanding of the clonal replacement phenomenon and update knowledge about MRSA clones. The work provided here provides a strong rationale for expanded studies at the national level with integrated clinical and genomic data to evaluate the policies and strategies for managing MRSA infections and reduce the impact of this pathogen on morbidity and mortality in the region.

## Acknowledgments

We gratefully acknowledge the Departamento de Laboratorios Clínicos, Escuela de Medicina, Pontificia Universidad Católica de Chile for the isolates included in this study.

## Supplementary material

**Supplementary Figure 1.**
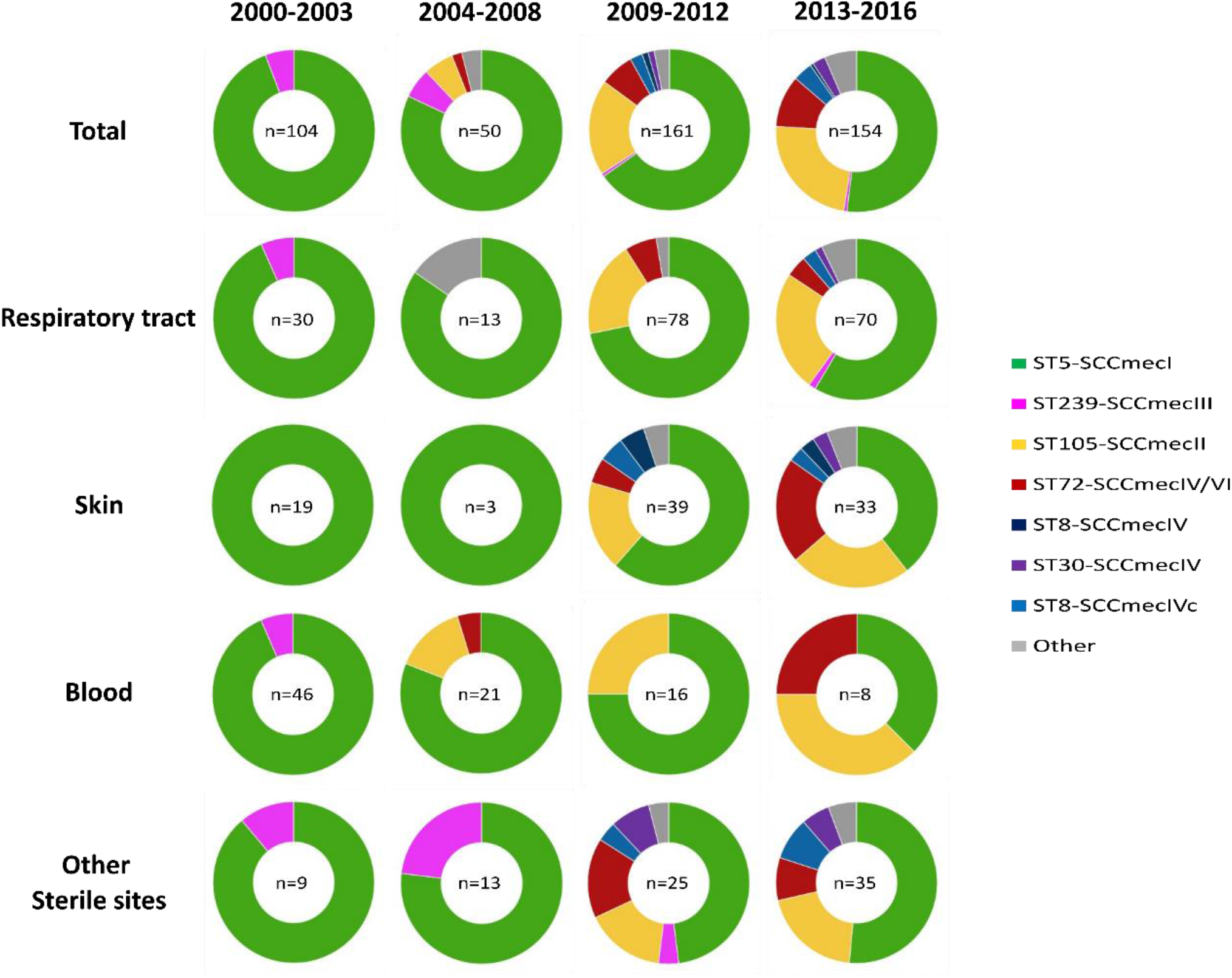
Temporal trends of the most frequent clones by anatomical sites.

**Supplementary table 1.** Antimicrobial resistance, virulence factors, and heavy metal resistance genes part of the core genome of the study collection.

| Gene | Type of function | Putative function |
| --- | --- | --- |
| *bla<sub>dha1</sub>* | Beta-lactam Resistance | Beta-lactamase, cephalosporinase |
| *AAC(3)* | Aminoglycoside Resistance | Acetylase |
| *APH(3')-III* | Aminoglycoside Resistance | Phosphotransferase |
| *norA* | Fluoroquinolone Resistance | Efflux pump |
| *norB* | Fluoroquinolone Resistance | Efflux pump |
| *mgrA* | Fluoroquinolone Resistance | Regulator of *norA*, *norB*, and *tet(38)* expression |
| *tet(38)* | Tetracycline Resistance | Efflux pump |
| *mepA* | Tetracycline Resistance | Efflux pump |
| *mepR* | Tetracycline Resistance | Regulator of *mepA* |
| *fosB* | Fosfomycin Resistance | A thiol transferase that leads to the resistance of fosfomycin |
| *lmrS* | Macrolide Resistance | Efflux pump |
| *hla* | Virulence | Alpha-hemolysin |
| *esxC* | Virulence | Type VII secretion system extracellular protein C |
| *essC* | Virulence | Type VII secretion system extracellular protein EssC |
| *esxB* | Virulence | Type VII secretion system extracellular protein B |
| *sspB* | Virulence | Cysteine protease that plays an important role in the inhibition of host innate immune response |
| *arsB* | Arsenic Resistance | Arsenical pump mambrane protein |
| *fieF* | Iron Resistance | Metal cation transporter |
| *czcD* | Cobalt and Cadmium Resistance | Metal cation efflux system protein |
| *copA* | Copper Tolerance | Copper exporting ATPase |

